# A simple and rapid method for isolating high-quality RNA from kenaf containing high polysaccharide and polyphenol contents

**DOI:** 10.1101/2020.07.06.189506

**Authors:** Xiaofang Liao, Hongwei Li, Aziz Khan, Yanhong Zhao, Wenhuan Hou, Xingfu Tang, Kashif Akhtar, Ruiyang Zhou

## Abstract

The isolation of high-quality RNA from kenaf is crucial for genetic and molecular biology studies. However, high levels of polysaccharide and polyphenol compounds in kenaf tissues could irreversibly bind to and coprecipitate with RNA, which complicates RNA extraction. In the present study, we proposed a simplified, time-saving and low-cost extraction method for isolating high quantities of high-quality RNA from several different kenaf tissues. RNA quality was measured for yield and purity, and the proposed protocol yielded high quantities of RNA (10.1-12.9 μg/g·FW). Spectrophotometric analysis showed that A_260_/_280_ ratios of RNA samples were in the range of 2.11 to 2.13, and A_260_/_230_ ratios were in the range of 2.04-2.24, indicating that the RNA samples were free of polyphenols, polysaccharides, and protein contaminants after isolation. The method of RNA extraction presented here was superior to the conventional CTAB method in terms of RNA isolation efficiency and was more sample-adaptable and cost-effective than commercial kits. Furthermore, to confirm downstream amenability, the high-quality RNA obtained from this method was successfully used for RT-PCR, real-time RT-PCR and Northern blot analysis. We provide an efficient and low-cost method for extracting high quantities of high-quality RNA from plants that are rich in polyphenols and polysaccharides, and this method was also validated for the isolation of high-quality RNA from other plants.

## 1. Background

Obtaining RNA of high purity and integrity is important for conducting analytical studies, such as reverse transcription, quantitative real-time polymerase chain reaction (qRT-PCR), Northern blot and complementary DNA (cDNA) library construction, in plant molecular biology. However, isolation of high-quality RNA from higher plant tissues is a challenging process due to the interference of endogenous RNase activation and external RNase introduction. In particular, tissues that are rich in polysaccharides, polyphenolic compounds and other types of secondary metabolites complicate RNA isolation (Tong et al., 2012). Polysaccharides are visually evident by their viscous, glue-like texture and make the pipetting of nucleic acids unmanageable (Wang and Stegemann, 2010), and they can also coprecipitate with nucleic acids and constitute the major hurdle for RNA isolation (Ding et al., 2008). The coprecipitation of these compounds with RNA reduces yield and increases the possibility of rapid degradation, making the sample unsuitable for further downstream applications due to the severe inhibition of enzymatic activity (Yang et al., 2017; Fang et al., 1992; Moser et al., 2004). In turn, polyphenols are known to be readily oxidized to form quinones that can irreversibly interact with proteins and nucleic acids to form high-molecular-weight complexes that hinder isolation of good-quality RNA (Japelaghi et al., 2011). Furthermore, with maturity or stressful plant growth conditions, tissues contain increased quantities of polyphenols and polysaccharides, which may further encumber the isolation of high-quality RNA (Choudhary et al., 2016). Hence, with the removal of such components is necessary to achieve the isolation of RNA of high quantity and quality. Many specific protocols, including those utilizing cetyltrimethylammonium bromide (CTAB)/NaCl (Chang et al., 1993), CTAB/LiCl (Khairul-Anuar et al., 2019), TRIzol (Guan et al., 2019) and sodium dodecyl sulfate (SDS) (Ma et al., 2015), designed for plants with a high content of polysaccharides and polyphenols, have been developed and used to render a high quantity of high-quality total RNA from young leaves (Dash, 2013). However, the majority of them pose certain limitations, as they are often expensive (Li et al., 2017), time consuming (Sivakumar et al., 2007), tissue specific (Guan et al., 2019) or technically complex (Japelaghi et al., 2011).

Kenaf (*Hibiscus cannabinus*) is an important fiber crop that is widely used in paper-making and weaving (Monti and Alexopoulou, 2013) and harbors significant heterosis in terms of phloem fiber production. Therefore, understanding the genetic factors underlying heterosis in kenaf holds promise for its breeding and production. Nucleic acid extraction from kenaf is notoriously difficult because kenaf has high concentrations of polysaccharides and polyphenols, which could coprecipitate with RNA and inhibit the enzymatic reactions in subsequent steps (Japelaghi et al., 2011). Several protocols, such as those of commercial kits, have been reported for RNA isolation from kenaf tissues (Li et al., 2017; Tang et al., 2019). However, the obtained RNA was contaminated by polysaccharides and easily degraded by residual RNase that could not be inhibited. Furthermore, although the extraction of nucleic acids was shortened using commercial reagent kits compared to conventional CTAB methods, these are not only expensive, but also contaminated by phenolic compounds, saccharides and proteins, resulting in low-quality RNA that is unsuitable for downstream applications.

Here, we present a simplified, efficient and cost-effective method for the isolation of high-quality total RNA from kenaf anthers, petals, leaves, stems and roots. For comparative purposes, we established a protocol for RNA extraction using CTAB in the extraction buffer, lithium chloride (LiCl) for precipitation (Zhou et al., 2015), and a commercially available ready-to-use reagent (Huayueyang, Beijing) for nucleic acid extraction. In contrast to the other tested methods, the RNA prepared by this protocol represented a significant improvement in terms of time savings and economics; additionally, the RNA was of high quality, extracted in high quantities and successfully used for downstream applications such as RT-PCR, qRT-PCR and Northern blot analysis. The proposed protocol was subsequently employed for successful RNA extraction from other plants, such as cotton, Roselle and soybean.

## 2. Materials and methods

### 2.1. Plant material

Kenaf was cultivated in the test field of Guangxi Academy of Agricultural Sciences (Nanning, Guangxi, China), and leaves, stems, roots, petals and anthers were collected at the flowering stage. Both cotton anthers and Roselle calyxes were collected at the flowering stage as well. Soybean stem were harvested from sixty-day-old plants grown in the field. All samples were flash-frozen in liquid nitrogen and stored at −80 °C for further use.

### 2.2. RNA extraction

#### (1) Preparation

Prior to RNA extraction, mortars and pestles were wrapped in tin foil and baked in an oven at 180 °C for 6 hours. All glassware was treated with 0.01% (v/v) diethylpyrocarbonate (DEPC) and autoclaved, and consumables (tips and tubes) were certified as RNase-free. All chemicals used were of molecular biology grade and purchased from Solarbio (Beijing, China). To limit exposure to noxious components (e.g., β-mercaptoethanol (B-ME), guanidine thiocyanate and phenol-chloroform), RNA extraction was conducted in a fume hood.

#### (2) Solutions and reagents

The RNA extraction buffer was composed of 100 mM Tris (pH 8.0), 25 mM EDTA (pH 8.0), 2 M NaCl, 2% CTAB (w/v) and 7% B-ME (V/V)^1^. Other solutions used in this study included a saturated guanidinium isothiocyanate solution (5 M), 100 and 75% (v/v) ethanol, RNase-free distilled deionized water, and water-saturated phenol: chloroform: isoamyl alcohol (PCI) 25:24:1 (v/v/v). All solutions were prepared with 0.01% DEPC-treated distilled water. RNA isolation was carried out twice using independent pools of tissue samples.

#### (3) Nucleic acid isolation procedure

Nucleic acid isolation was carried out using the following steps:

➀ Initially, approximately 0.2-0.5 g of plant tissue sample was ground into a fine powder using a prechilled mortar and pestle under liquid nitrogen. Subsequently, the homogenized sample was immediately transferred into a 2-mL RNase-free centrifuge tube to minimize any RNA degradation.
➁ A total of 1 mL of extraction buffer that had been preheated to 65 °C was added, and the suspension was thoroughly mixed and incubated for 10 min at 65 °C for adequate lysis.
➂ An equal volume of PCI (25: 24:1 v/v/v) was added and mixed thoroughly, and the mixture then centrifuged at 18,000 ×g for 10 min at 4 °C.
➃ The aqueous phase was transferred into a new 2-mL RNase-free tube, and saturated guanidine isothiocyanate (5 M) was added at a ratio of 1.7/1 according to the volume of supernatant. Then, 0.5-fold volume anhydrous ethanol was added and mixed gently.
➄ The mixture was transferred to an RNA purification column and centrifuged at 18,000 ×g for 30 s at 4 °C.
➅ The eluate in the collection tube was discarded, and step ➄ was repeated until all the mixture had been processed.
➆ A total of 750 μL of ethanol (75%) was added to the RNA spin column, which was then centrifuged at 18,000 ×g for 30 s at 4 °C to wash the spin column membrane. The eluate in the collection tube was discarded.
➇ The RNA spin column was centrifuged for 2 min at full speed and 4 °C to ensure that no ethanol remained in the column.
➈ The RNA spin column was placed in a new 1.5-mL RNase-free tube. Then, 50 μL of RNase-free deionized water was added directly to the spin column membrane, and the column was placed on ice for 2 min. The tube was centrifuged for 2 min at 18,000 × g at 4 °C to elute the RNA.

The other two comparative RNA extraction methods included the protocol described by Zhou (2015) and a ready-to-use RNA extraction kit (Huayueyang, Beijing) that was implemented following the manufacturer’s instructions. Each kenaf tissue type (anthers, petals, mature leaves, roots and stems) was extracted in two independent experiments and measured by each method.

### 2.3. Assessment of RNA quantity and quality

The purity and concentration of our RNA extracts were determined using a spectrophotometer (NanoDrop 2000, Thermo Scientific, USA) by measuring absorbance ratios of A_260_/A_280_ and A_260_/A_230_. The integrity of total RNA was verified by running a 500-ng sample in a 1% (w/v) agarose gel that was then stained with SYBR Green II (Tiandz, China) and imaged using a UV transilluminator from Syngene (Bio-Rad, USA).

### 2.4. Reverse Transcriptase polymerase chain reaction (RT-PCR), real-time RT-PCR and Northern blots

The isolated RNA samples were reverse transcribed to confirm downstream amenability. One microgram of total RNA was used in a reverse transcription reaction using TransScript One-Step gDNA removal and cDNA synthesis SuperMix (TransGene Biotech, Beijing, China) to obtain 20 μL of cDNA solution following the manufacturer’s instructions. The cDNA was PCR amplified by *cox2*, in which primers were designed to span introns and amplify a 717-bp fragment for cDNA and a 2208-bp fragment for gDNA, including a 1491-bp intron (primers are shown in Table 1). PCRs were carried out in a final volume of 20 μL of reaction mixture containing 10 μL of 2 × Taq Master Mix (Vazyme, China), 0.3 μM each forward and reverse primer and 50 ng of cDNA template. The thermal cycling program for PCR amplification was as follows: predenaturation at 94 °C for 3 min, followed by 35 cycles of 40 s at 94 °C for denaturation, 1 min at 58 °C for annealing, 2 min at 72 °C for extension, and a final step of 5 min at 72 °C. The amplified product was visualized by gel electrophoresis in 1% (w/v) agarose gel. We also presented some samples for the expression of the mitochondrial genes *cox3* and *atp9* using real-time RT-PCR and Northern blot analysis following the procedure described by Liao et al (2016). All primer sequences used in this study are listed in Table 1.

**Table 1.**
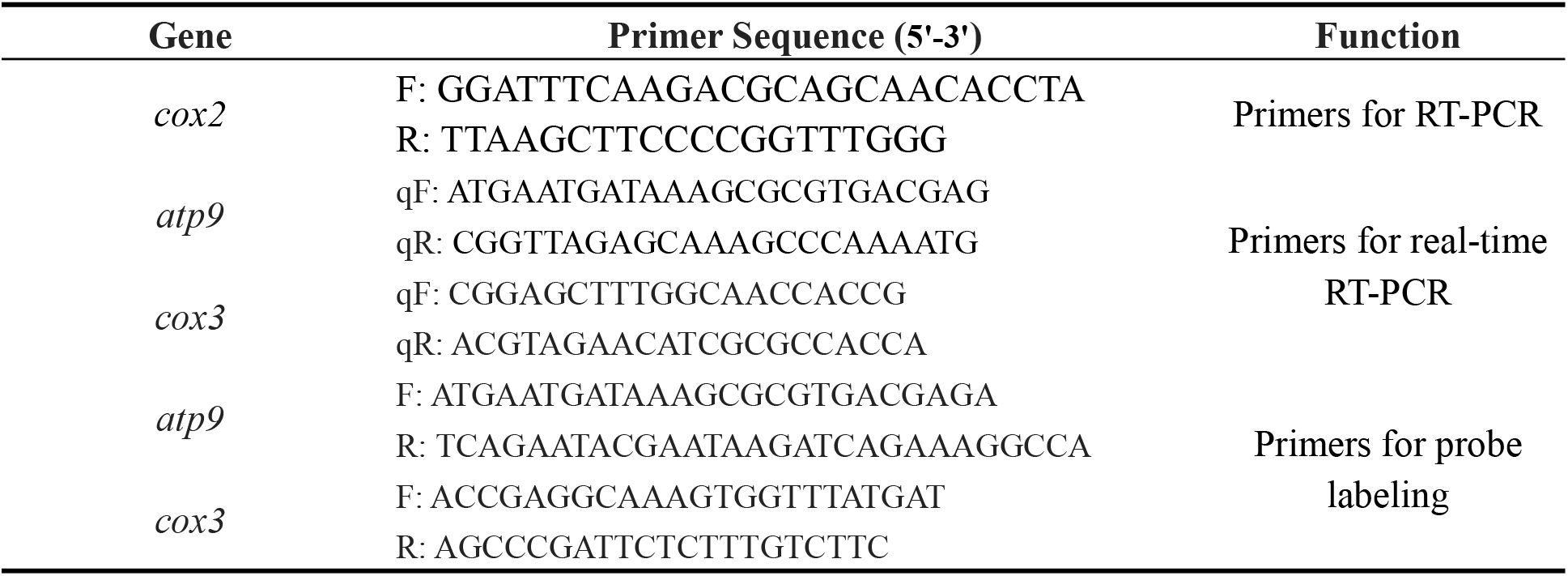
Primers used for this study.

## 3. Results and discussion

### 3.1. Quantity, quality and yield of total RNA

Several total RNA extraction protocols specifically designed for kenaf have been reported. However, these protocols present important limitations: many reported methods utilize expensive commercial extraction kits (Li et al., 2017; Tang et al., 2019), others were time consuming, which could cause a potential risk of RNA degradation in the process of extraction (Zhou et al., 2015), while other kenaf-based protocols are for specific tissues (Chen et al., 2011). In the current study, we described a rapid, efficient and reliable protocol that allowed for the extraction of high-quality total RNA from several kenaf tissues. With our method, we successfully isolated high-quality RNA from kenaf anthers (Fig. 1: lanes 1-2), sepals (Fig. 1: lanes 3-4), mature leaves (Fig. 1: lanes 5-6), roots (Fig. 1: lanes 7-8) and stems (Fig. 1: lanes 9-10). RNA samples showed two sharp and well-resolved ribosomal bands corresponding to the 28S and 18S rRNAs on 1.0% agarose gels (Fig. 1), and the 28S rRNA band was twice as abundant as the 18S rRNA band (Fig. 1), indicating that little or no RNA samples isolated from the different tissues were degraded during the extraction. Spectrophotometric analysis revealed A_260_/_A280_ ratios ranging between 2.11 and 2.15 (Table 1), indicating a lack of protein contamination. Similarly, the A_260_/A_230_ ratios of all tested samples were greater than 2.0, indicating that the obtained total RNA was highly pure and free of protein, polyphenol and polysaccharide contamination (Dash, 2013; Guan et al., 2019; Khairul-Anuar et al., 2019). The yields of total RNA extracted from different kenaf tissues were quite diverse depending on the tissue sources. The kenaf anthers yielded the highest amount of total RNA (229.31 μg/g fresh weight (FW)), and the root issues displayed the lowest yield levels (76.24 μg/g FW) (Table 1), indicative of the higher number of RNA cells in kenaf anthers than in roots, resulting in high sample recovery.

**Fig. 1.**
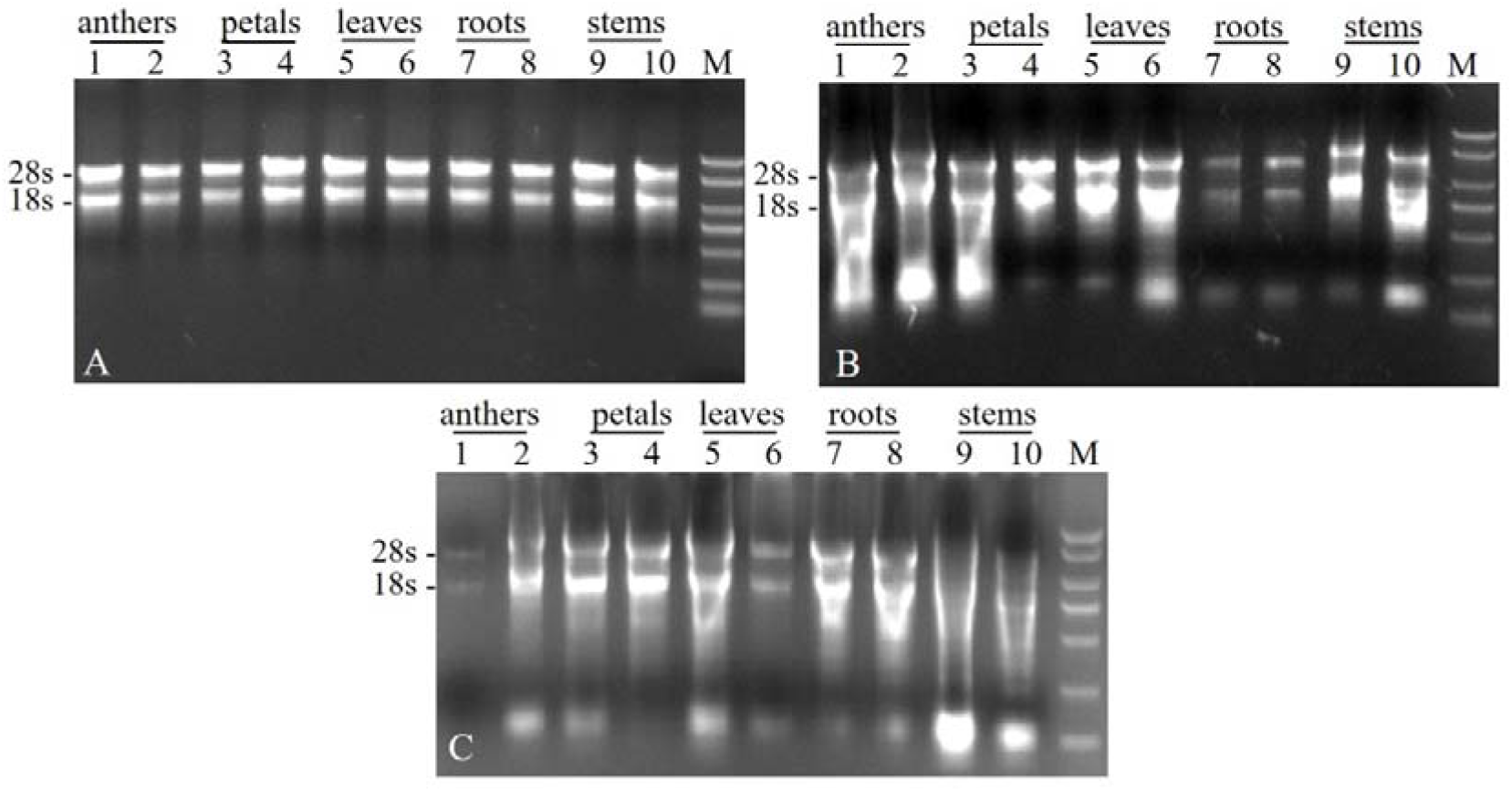
Gel electrophoresis of kenaf RNA isolated from different tissues using the proposed protocol (A), an RNA extraction kit (B) and a Li-Cl-based method (Zhou et al., 2015) (C). M, BM 5000 marker.

### 3.2. Comparison with the other RNA extraction methods

Nucleic acids isolated from kenaf tissues following the proposed protocol were compared in terms of quantity and quality with those isolated from the LiCl-based protocol proposed by Zhou (2015) and the ready-to-use RNA extraction kit (Huayueyang, Beijing). The quantity and purity of kenaf RNA preparations when using this simple protocol were found to be superior to those obtained using the previously reported protocols (Table 2 and Fig. 1). As shown in Table 2, the total nucleic acid quantity was the highest when using the current protocol according to spectrophotometric quantification, followed by similar amounts when using the RNA extraction kit, and the yield was the lowest when using the method described by Zhou (2015). However, gel electrophoresis showed that nucleic acids extracted from kenaf tissues were nearly degraded, especially when using the Li-Cl-based protocol described by Zhou (2015) (Fig. 1B and 1C). Nucleic acid purity indicated by A_260_/A_280_ absorbance ratios was between 1.48 and 1.94 for the LiCl-based and RNA extraction kit protocols, respectively. This suggested that the high yields shown by spectrophotometric analysis were likely the result of false measurements of secondary metabolite contaminants (Wang et al., 2012), as polysaccharides and polyphenolic compounds often coprecipitate and contaminate nucleic acids during extraction, thereby affecting both the quality and quantity of isolated nucleic acids (Kansal et al., 2008). Similarly, A_260_/A_230_ absorbance ratios were lower (ranging between 0.55 and 1.41) following these two protocols, indicative of contamination with polyphenols and polysaccharides (Kim and Hamada, 2005).

**Table 2.**
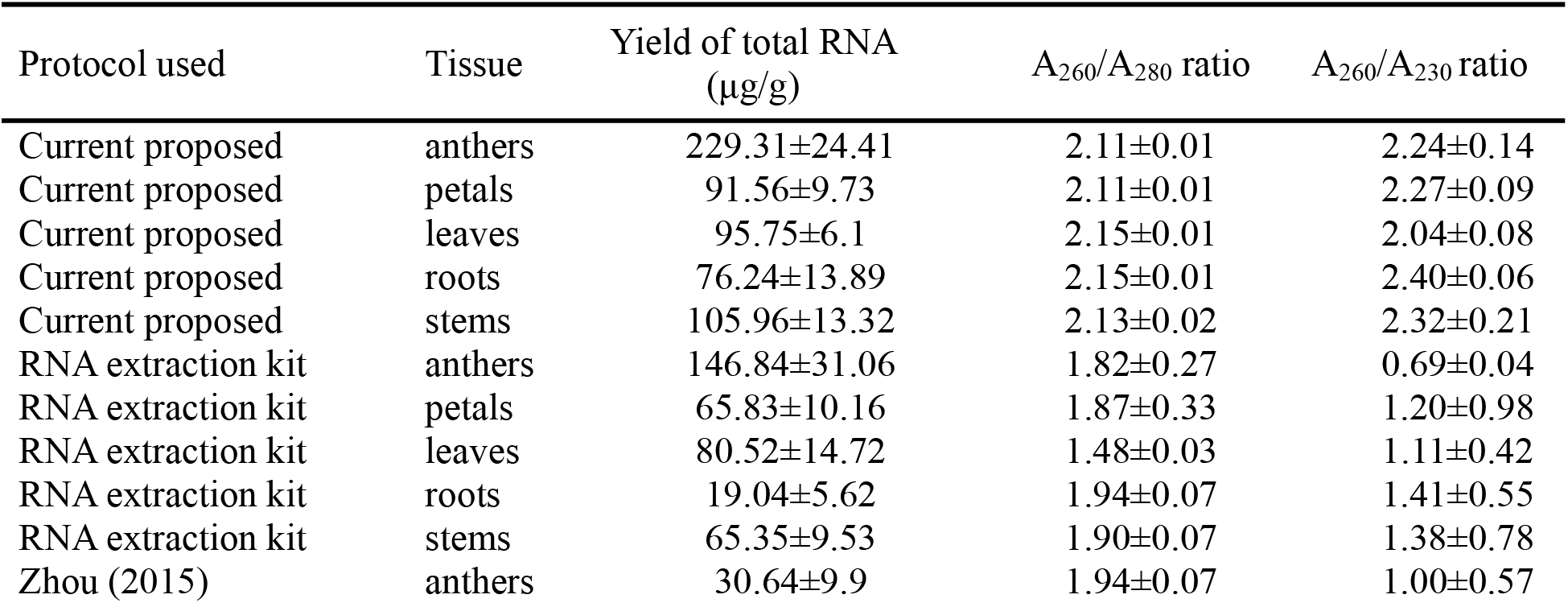
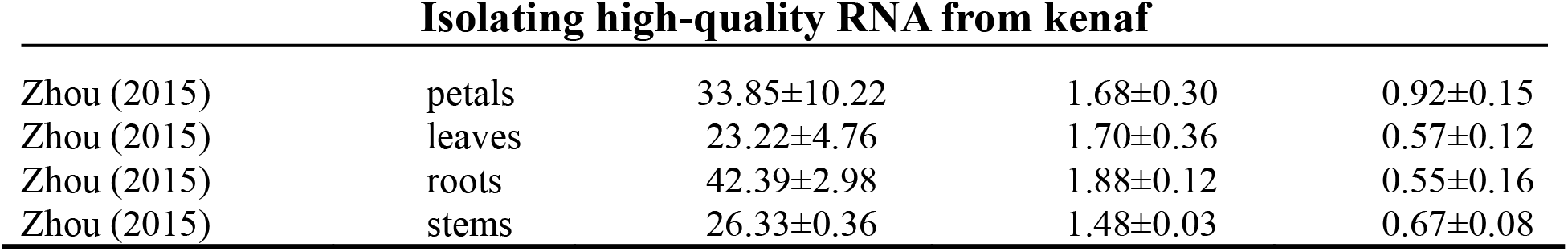
The purity and quality determination of RNA from kenaf tissues isolated by three different protocols.

| Protocol used | Tissue | Yield of total RNA<br>($\mu\text{g/g}$) | $A_{260}/A_{280}$ ratio | $A_{260}/A_{230}$ ratio |
| --- | --- | --- | --- | --- |
| Current proposed | anthers | $229.31 \pm 24.41$ | $2.11 \pm 0.01$ | $2.24 \pm 0.14$ |
| Current proposed | petals | $91.56 \pm 9.73$ | $2.11 \pm 0.01$ | $2.27 \pm 0.09$ |
| Current proposed | leaves | $95.75 \pm 6.1$ | $2.15 \pm 0.01$ | $2.04 \pm 0.08$ |
| Current proposed | roots | $76.24 \pm 13.89$ | $2.15 \pm 0.01$ | $2.40 \pm 0.06$ |
| Current proposed | stems | $105.96 \pm 13.32$ | $2.13 \pm 0.02$ | $2.32 \pm 0.21$ |
| RNA extraction kit | anthers | $146.84 \pm 31.06$ | $1.82 \pm 0.27$ | $0.69 \pm 0.04$ |
| RNA extraction kit | petals | $65.83 \pm 10.16$ | $1.87 \pm 0.33$ | $1.20 \pm 0.98$ |
| RNA extraction kit | leaves | $80.52 \pm 14.72$ | $1.48 \pm 0.03$ | $1.11 \pm 0.42$ |
| RNA extraction kit | roots | $19.04 \pm 5.62$ | $1.94 \pm 0.07$ | $1.41 \pm 0.55$ |
| RNA extraction kit | stems | $65.35 \pm 9.53$ | $1.90 \pm 0.07$ | $1.38 \pm 0.78$ |
| Zhou (2015) | anthers | $30.64 \pm 9.9$ | $1.94 \pm 0.07$ | $1.00 \pm 0.57$ |
| Zhou (2015) | petals | 33.85±10.22 | 1.68±0.30 | 0.92±0.15 |
| Zhou (2015) | leaves | 23.22±4.76 | 1.70±0.36 | 0.57±0.12 |
| Zhou (2015) | roots | 42.39±2.98 | 1.88±0.12 | 0.55±0.16 |
| Zhou (2015) | stems | 26.33±0.36 | 1.48±0.03 | 0.67±0.08 |

In the proposed method, CTAB, a strong ionic denaturing detergent, at a final concentration of 2% (w/v) and a high concentration of B-ME (7% (v/v)) were added in the extraction buffer. This could completely solubilize the cell membranes and bind to the nucleic acid, as well as eliminate most of the polysaccharides and polyphenolic compounds (Kim and Hamada, 2005; Wang et al., 2008). In addition, a relatively high NaCl content (2 M) in the extraction buffer promoted salting out of the protein and avoided coprecipitation of polysaccharides with the RNA while leaving the RNA in solution (El-Ashram et al., 2016). Moreover, guanidinium isothiocyanate is an RNase inhibitor that can effectively inhibit the activity of RNase during extraction (Suzuki et al., 2001).

Furthermore, the presence of precooled 5 M guanidinium isothiocyanate and absolute ethyl alcohol that could bind to nucleic acids formed a jelly-like precipitate that could then be adsorbed by an RNA purification column. Taken together, these results show that good-quality total RNA was successfully obtained from kenaf tissues with high levels of secondary metabolites. The current protocol of RNA isolation was not only efficient but also required less time (~ 1.5 h) that earlier methods described by Chen et al (2011) and Zhou et al (2015), which required ~ 8 h. Simultaneously, lower amounts of reagents were used throughout the current procedure, thus contributing to reduced costs.

### 3.3. Reverse transcription and downstream application of the RNA

The RNA isolated by the proposed method was tested by amplifying the *cox2* gene fragment. A 2208-bp fragment was amplified using genomic DNA as a template, and a 717-bp fragment was amplified using cDNA as a template (Fig. 2). This suggested that the genomic DNA was successfully eliminated from all samples by the gDNA remover reagent during reverse transcription, and PCR products using cDNA as the template indicated high-quality RNA extraction. Furthermore, this differential amplification of fragments in genomic and cDNA samples not only proved the quality of isolated RNA samples and the absence of DNA contamination but also confirmed the amenability of this method for downstream processing without any ambiguity. Moreover, the isolation of RNA with the current protocol was successfully employed for real-time RT-PCR (qRT-PCR) gene expression analysis and Northern blot analysis. As shown in Fig. 3, the values of qRT-PCR cycle thresholds (Ct) ranged from 17 to 25 cycles (Fig. 3A and 3B), and the melting curve was specific, with a solitary peak occurring at approximately 82 °C to 83 °C (Fig. 3C-F). In addition to qRT-PCR, Northern blot analysis was carried out to further demonstrate the quality of RNA prepared by our protocol. The fluorescent signal with clear bands and high intensity was detected in RNA transcripts (Fig. 4), indicating that high-quality RNA was isolated from kenaf tissues and was suitable for downstream application.

**Fig. 2.**
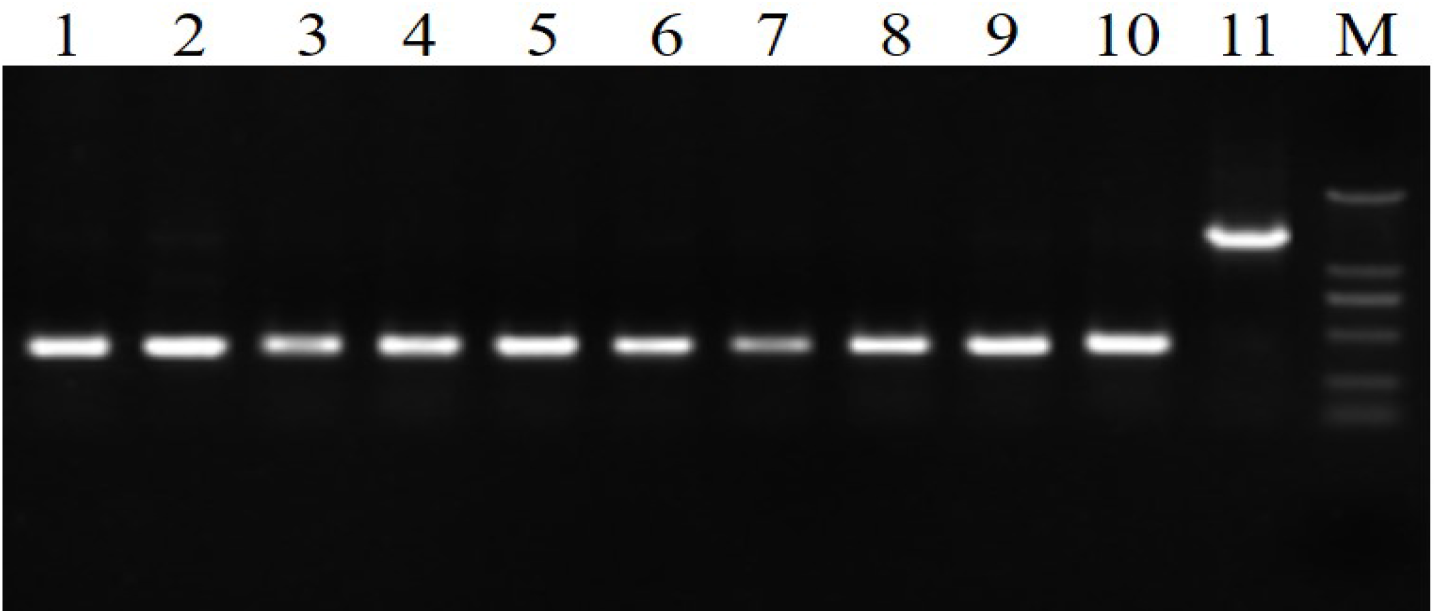
RT-PCR determination of the extracted RNA isolated by the current protocol. Lanes 1-10, the amplification products of RT-PCR that used cDNA from kenaf anthers, petals, leaves, roots and stems as templates. Lane 11, the PCR amplification product that used kenaf anther gDNA as the template. M, BM 5000 marker.

**Fig. 3.**
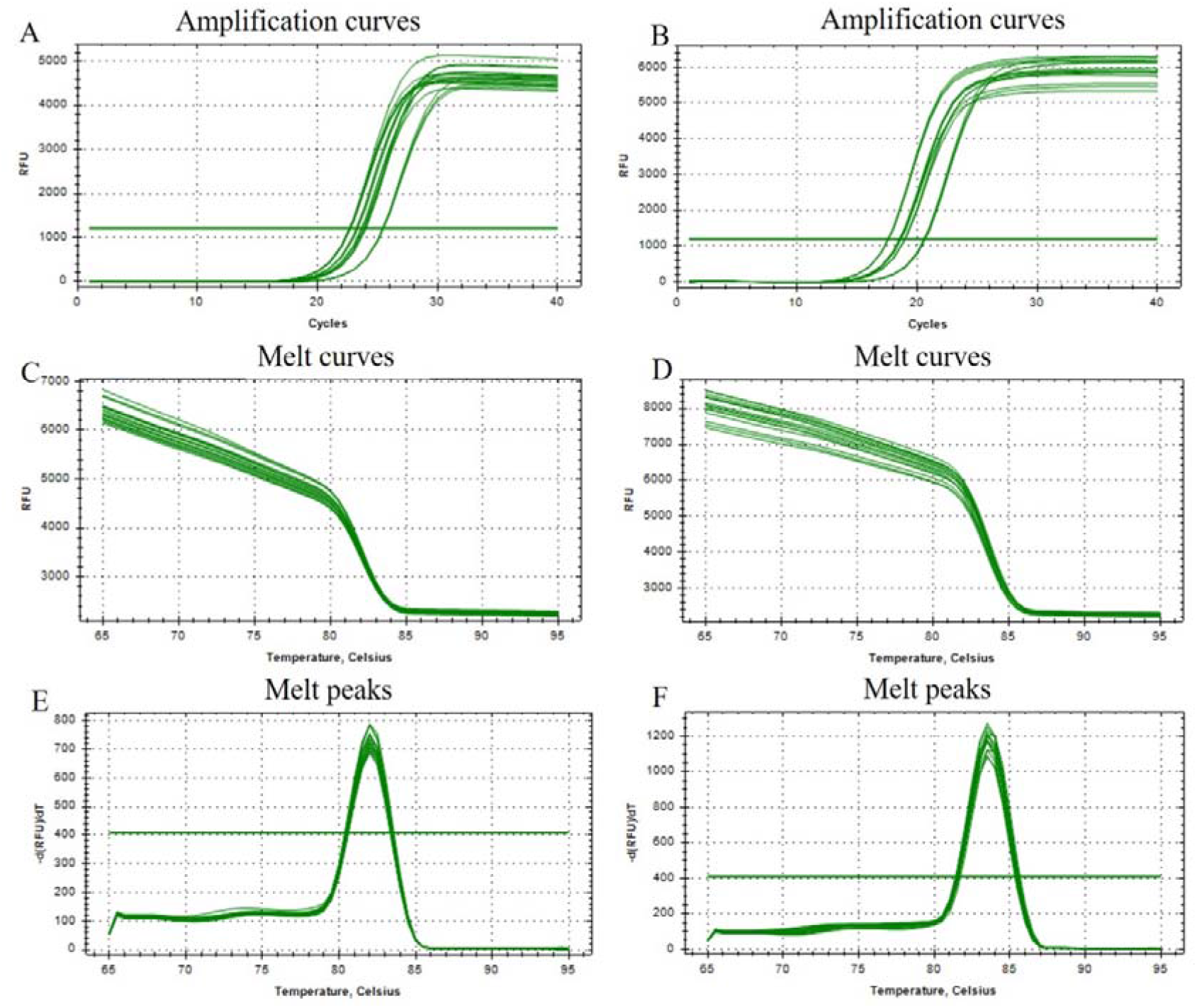
qPCR analysis of *cox3* and *atp9* genes in kenaf tissues. A-B, application curves of the qPCR products for *cox3* (left) and *atp9* (right); C-D, melting curves of the qPCR products for *cox3* (left) and *atp9* (right); E-F, melting peaks of the qPCR products for *cox3* (left) and *atp9* (right).

**Fig. 4.**
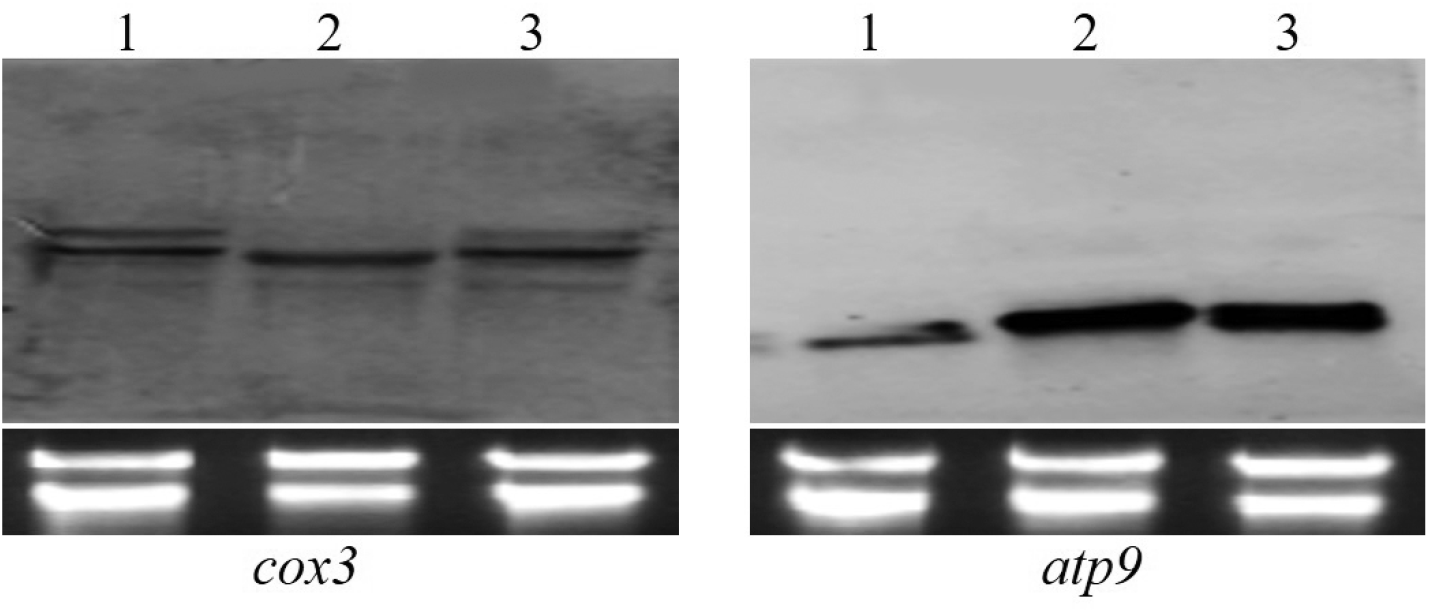
Northern blot analysis for *cox3* and *atp9* of three varieties of kenaf RNA extracted by the current protocol. The three varieties of kenaf shown in lanes 1-3 were UG93A, UG93B and F1 (UG93A/UG93R), respectively.

The present protocol could also be successfully applied for RNA extraction from other recalcitrant plant tissues, such as cotton and Roselle. The consistency of the results obtained from independent biological replicates confirmed the reproducibility of this method and provided a useful tool for molecular studies focusing on crops rich in secondary metabolites. The protocol was also successfully employed using soybean stems as material (Fig. 5) therefore potentially having a very wide range of applicability.

**Fig. 5.**
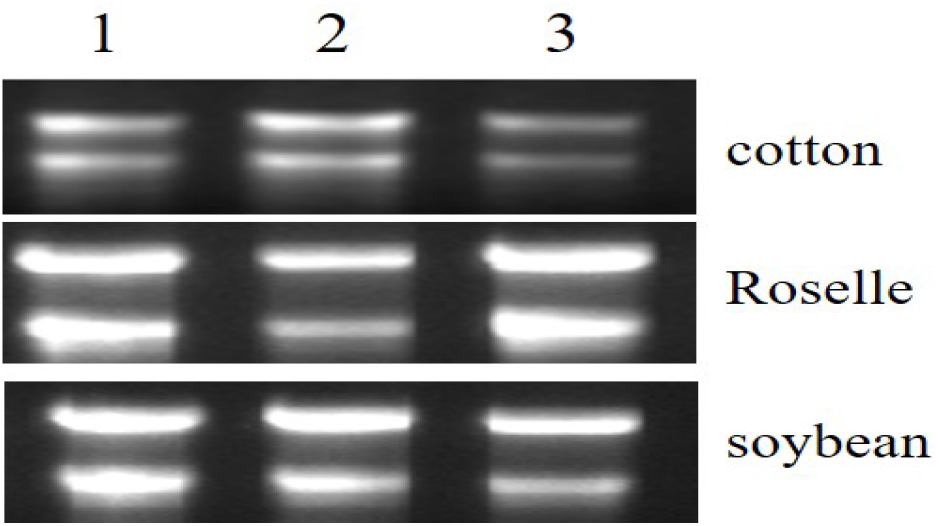
Gel electrophoresis of RNA isolated from different plants using the proposed protocol. Lanes 1-3 represent three independent biological replicates.

## 4. Conclusions

We reported a new protocol for the extraction of high quantities of high-quality RNA from kenaf tissues with high levels of polysaccharides and polyphenols, and the extracted RNA was suitable for subsequent gene isolation and expression experiments, such as reverse transcription, qRT-PCR, and Northern blot analysis. This protocol is an easy, efficient, relatively inexpensive, and highly reproducible method to isolate RNA from kenaf, especially from tissues such as anthers, petals and leaves, which are rich in polyphenols and polysaccharides. Thus, this simple and fast protocol has been routinely used in our laboratory for RNA isolation from cotton, Roselle, soybean stems and other plants with high levels of secondary metabolites. This protocol greatly reduces labor time and costs without compromising the quality and yield of RNA samples.

## Acknowledgments

This study was supported by the National Science Foundation of China (No. 31571719 and No. 31660430), the Natural Science Foundation of Guangxi Province (No. 2018JJB130045 and 2019JJA130200), and Basic Business Expenses Project of Guangxi Academy of Agricultural Sciences (No. Guinongke2019M23).

## Author Contributions

R. Z. initiated the experiment. X. L. conducted the experiment and drafted the manuscript. H. L. isolated the RNA. Y. Z. and B. Z. conducted the Northern blot analysis. W. H. and X. T. assisted with the experiment. C. L. provided suggestions for the manuscript. A. K. and K. A. revised the manuscript and edit language.

## Footnotes

1 Notes: added just before use.

